# Hype versus hope: Deep learning encodes more predictive and robust brain imaging representations than standard machine learning

**DOI:** 10.1101/2020.04.14.041582

**Authors:** Anees Abrol, Zening Fu, Mustafa Salman, Rogers Silva, Yuhui Du, Sergey Plis, Vince Calhoun

## Abstract

Previous successes of deep learning (DL) approaches on several complex tasks have hugely inflated expectations of their power to learn subtle properties of complex brain imaging data, and scale to large datasets. Perhaps as a reaction to this inflation, recent critical commentaries unfavorably compare DL with standard machine learning (SML) approaches for the analysis of brain imaging data. Yet, their conclusions are based on pre-engineered features which deprives DL of its main advantage: representation learning. Here we evaluate this and show the importance of representation learning for DL performance on brain imaging data. We report our findings from a large-scale systematic comparison of SML approaches versus DL profiled in a ten-way age and gender-based classification task on 12,314 structural MRI images. Results show that DL methods, if implemented and trained following the prevalent DL practices, have the potential to substantially improve compared to SML approaches. We also show that DL approaches scale particularly well presenting a lower asymptotic complexity in relative computational time, despite being more complex. Our analysis reveals that the performance improvement saturates as the training sample size grows, but shows significantly higher performance throughout. We also show evidence that the superior performance of DL is primarily due to the excellent representation learning capabilities and that SML methods can perform equally well when operating on representations produced by the trained DL models. Finally, we demonstrate that DL embeddings span a comprehensible projection spectrum and that DL consistently localizes discriminative brain biomarkers, providing an example of the robustness of prediction relevance estimates. Our findings highlight the presence of non-linearities in brain imaging data that DL frameworks can exploit to generate superior predictive representations for characterizing the human brain, even with currently available data sizes.

## Introduction

The application of machine learning for investigation of neurological and psychiatric disorders has grown greatly in the last two decades ^1,2^. Standard machine learning (SML) approaches predict health-related outcomes by manipulating specific linear or non-linear prediction functions using rules of inference. An indispensable prerequisite to boosting the performance of SML approaches is reducing the dimensionality of the input space, typically enabled through hand-crafted or expert-designed feature selection (i.e. identification of a subset of variables that capture most of the information in the data) and/or feature extraction (i.e. projection of the features onto a lower-dimensional space by some linear or non-linear data transformation) techniques ^3,4^. The persistent challenge imposed by this preliminary step paved the way for the introduction of deep learning (DL) approaches. The DL approaches instead, can exploit the wealth of information available from minimally preprocessed input images to characterize the subtle patterns inherent in the input data as an integral part of the training process. The training phase of DL approaches often involves the automatic and adaptive discovery of discriminative data representations at multiple levels of hierarchy in an end-to-end (input to output) learning procedure. Application of this radically different approach in an end-to-end manner provisions backwards mapping to the input image space through methodical interpretations, thus allowing one to delineate the features in the input space that are most influential in predicting an attempted task. On the contrary, relevant spatial relationships may be lost at the dimensionality reduction stage, arguably, required for SML methods to work. DL approaches have been successful in learning more predictive data encodings (i.e. representations) compared to their manually engineered counterparts and several automated dimensionality reduction techniques in the field of computer vision ^5,6^.

DL approaches have already shown great promise in diverse applications to medical imaging data ^7-10^. Several structural and functional brain imaging modalities are now being actively used to study mental health non-invasively. While the SML approaches have contributed to these efforts making significant advances ^11-16^, the relatively newer whole-brain DL approaches are just beginning to record successes, particularly in the image preprocessing, diagnostic classification, regression, disease characterization, and disease prediction domains ^17-24^. Concurrently, similar to preceding influential technologies, expectations of future performance of DL frameworks sometimes grow out of proportion. For example, their viability to learn subtle properties of complex multiscale brain imaging data and potential to scale may be hyped ^25^. Perhaps as a reaction to this inflation, recent critical commentaries unfavorably compare DL with the SML approaches ^26,27^. Yet, the conclusions in these commentaries are based on analyses which use pre-engineered features, hence depriving DL of its main advantage: representation learning.

In this work, we correct for deficiencies of prior benchmarks in a principled, comparative analysis of structural brain imaging data and show how vital the representation learning part of DL is for its performance. To this end, we systematically profile the classification performance and empirical time complexity of several SML and DL methods on a ten-way age and gender-based classification task ^26^ using a large dataset of structural magnetic resonance imaging (sMRI) images. We conduct this comparison for a range of training sample sizes to compare and contrast the asymptotic behavior in performance improvement and relative time complexity of the two approaches. Additionally, we probe the consistency in the prediction power of the DL embeddings by evaluating the performance of SML methods trained on these features. Finally, we conduct post-hoc analyses to assess the degree of consistency and robustness of the validated DL models. While the results from our study show the best performing approach for fitting a discriminative classifier for the attempted sample task and brain imaging modality, a similar analysis could easily be extended to other inference tasks and modalities. Next, we present the findings of our exploratory work.

## Results

### DL Capacitates more Predictive Features

We systematically evaluated how the performance (as measured by accuracy and run time) of the SML and DL models scaled as a function of training sample size in a 10 class age and gender classification task (i.e. five age groups from each gender) evaluated in a standard repeated (*n* = 20), stratified cross-validation procedure as outlined in **Figure 1**. We used gray matter volume maps extracted from the structural MRI (sMRI) data of 12,314 unaffected (i.e. with no diagnosed or self-reported mental illness) subjects for this assessment. To establish a performance baseline for the SML methods, we included three linear SML models - linear discriminant analysis (LDA), logistic regression (LR) and support vector machine with a linear kernel (SVML) and three non-linear SML models - support vector machines with a polynomial (SVMP), radial-basis function (SVMR), and sigmoidal (SVMS) kernels (similar to Schulz, et al. ^26^). In addition, we tested two non-linear DL models - both 3D CNN variants of the AlexNet architecture ^28^ that differed primarily in the network depth (*depth*DL2 > *depth*DL1) and the number of channels in the convolutional layers. Given that feature extraction is an indispensable measure of boosting performance of linear and kernel based SML methods, we reduced the gray matter maps with three dimensionality reduction methods: Gaussian Random Projection (GRP), Recursive Feature Elimination (RFE) and Univariate Feature Selection (UFS) as detailed in the methods section. We trained the DL architectures directly on the unreduced input space of 3D gray matter maps to fully utilize their representational power.

**Figure 1:**
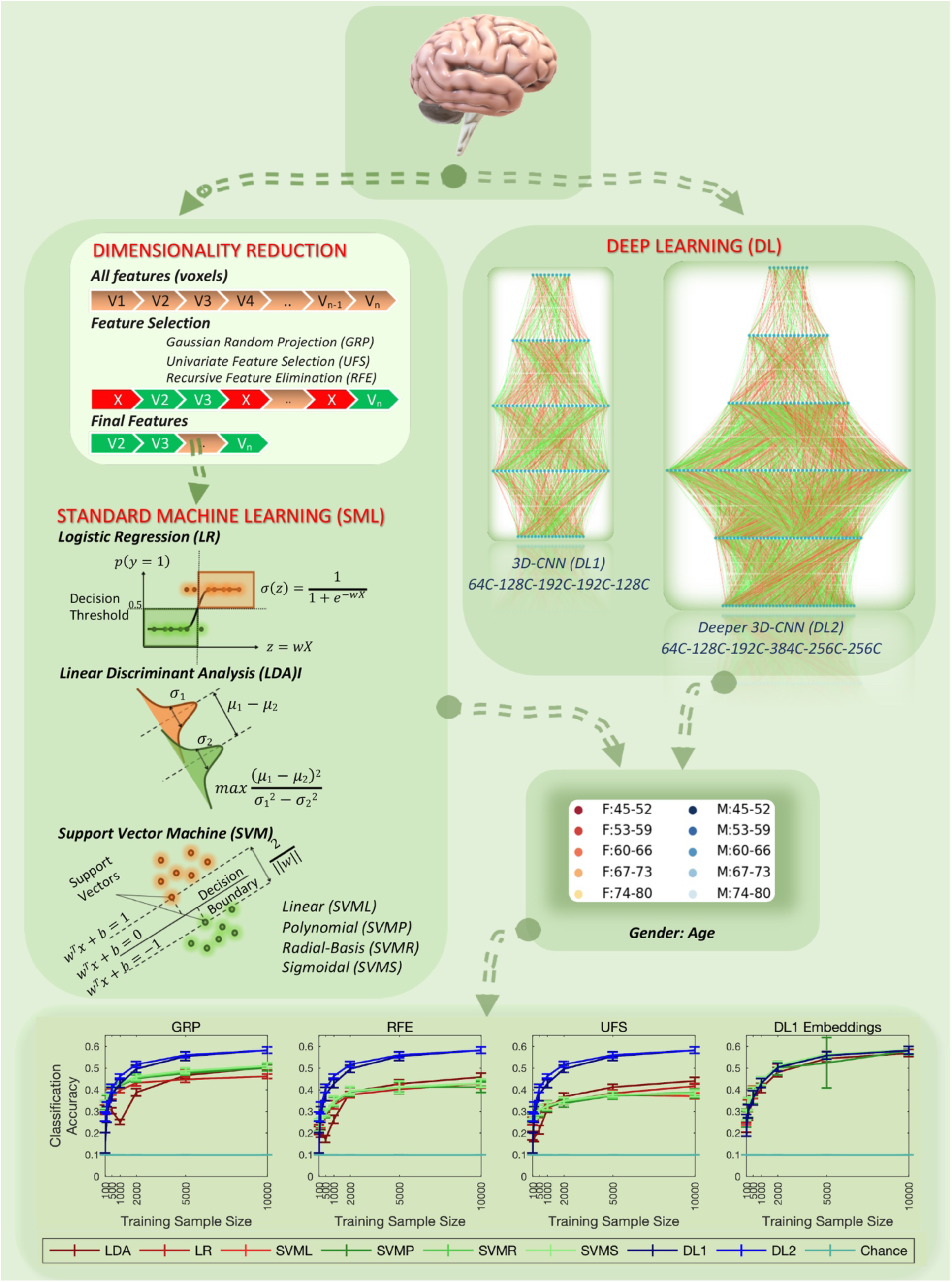
Systematic Comparison of Classification Performance on the 10-class Age and Gender Classification Task. The accuracy of all models was assessed in a 10 class classification task of a structural MRI data of 12,314 subjects via 20 random partitions of the data into training, validation, and test sets. For each repeat, the hyperparameters were tuned on the validation data, and reported performance was evaluated on held-out test set. The SML models were trained/tested on gray matter features reduced by three different feature reduction methods. In contrast, the DL models were trained on the unreduced (i.e. 3D voxel space) preprocessed gray matter maps. The tested DL models significantly (*p* < 0.005) outperformed all SML models regardless of the dimensionality reduction method. Finally, superior feature extraction of the DL models was immediately evident as the SML models trained on the DL representations performed equally well.

We found that the two DL models significantly outperformed the SML models evaluated on each of the reduced feature spaces (**Figure 1**). For the highest sample size (*n* = 10000), the DL models reported 10 class classification accuracies of 58.19% (D1) and 58.22% (D2) respectively (note, this task had a chance probability of 10%). In contrast, the SVMS and LDA models reported the highest accuracies for the GRP (SVMS: 51.15%), RFE (LDA: 45.77%) and UFS (LDA: 44.07%) features. Indeed, the GRP method resulted in the most predictive features for all SML models, followed by the RFE method. While both DL models consistently reported significant improvement (*p* < 0.005) with an increase in training sample size, this observation was not necessarily true for the SML models. For example, the performance of LDA on GRP features dropped initially, possibly due to a smaller training sample size than the validation and test data sizes. Additionally, as expected for sparse models, no significant improvement was observed in the performance of the SVMP for the RFE features, and SVML and SVMS for UFS features with an increase in training sample size from n = 5000 to n = 10000.

Interestingly, the performance improvement for DL models showed asymptotic behavior similar to SML methods, though with significantly higher performance. That is, for the DL methods as well, the performance gains are slowing down, although the models do continue to improve. Whether the slow down is impactful and where is the point of diminishing returns occurs is application dependent. Our observation demands a further confirmation as there are many ways to potentially score further gains in performance by testing even deeper models, finetuning the existing DL models, and exploring other DL approaches. Furthermore, if the DL models are indeed extracting superior (i.e. more predictive) features consistently, the lower dimensional encodings generated by them should result in significantly improved performance if used as input features of the SML models as compared to the three tested dimensionality reduction methods. To ascertain this, we conducted a post hoc analysis where we evaluated the performance of the SML models on the trained encodings from the DL1 model (i.e. the output of the first fully connected layer in DL1). As expected, we observed a significant increase in the performance of the SML methods applied to test data, providing evidence that the SML methods could perform equally well if using the DL encoded feature spaces (**Figure 1**).

### DL Presents Lower Empirical Asymptotic Complexity in Relative Computational Time

While DL models are notorious for high computational complexity, the high empirical computational requirements of the standard, CPU-based SML implementations on large training datasets are often overlooked. The fact that the computational complexity of most of the tested SML models is either cubic or quadratic in the number of input samples makes a comparison of empirical asymptotic time complexity between the two classes of methods on large datasets even more interesting. Hence, we pursued empirical evidence to determine the growth in the computational time of the two classes of methods as a function of training sample size. **Figure 2A** presents the average computation time of all tested models. This comparison illuminates a higher growth rate of computation time for most of the SML models, as the recorded differences for the two classes of models diminished with increasing training sample sizes for all SML models except LDA. Furthermore, to confirm if this observation indeed implied lower empirical asymptotic complexity for DL models, we estimated a *relative* computational growth rate metric by normalizing the computation time with the computation time for the smallest training sample size. The results of this analysis (**Figure 2B)** are an empirical evidence of lower growth rate in computational complexity of DL models compared to all SML models except LDA.

**Figure 2:**
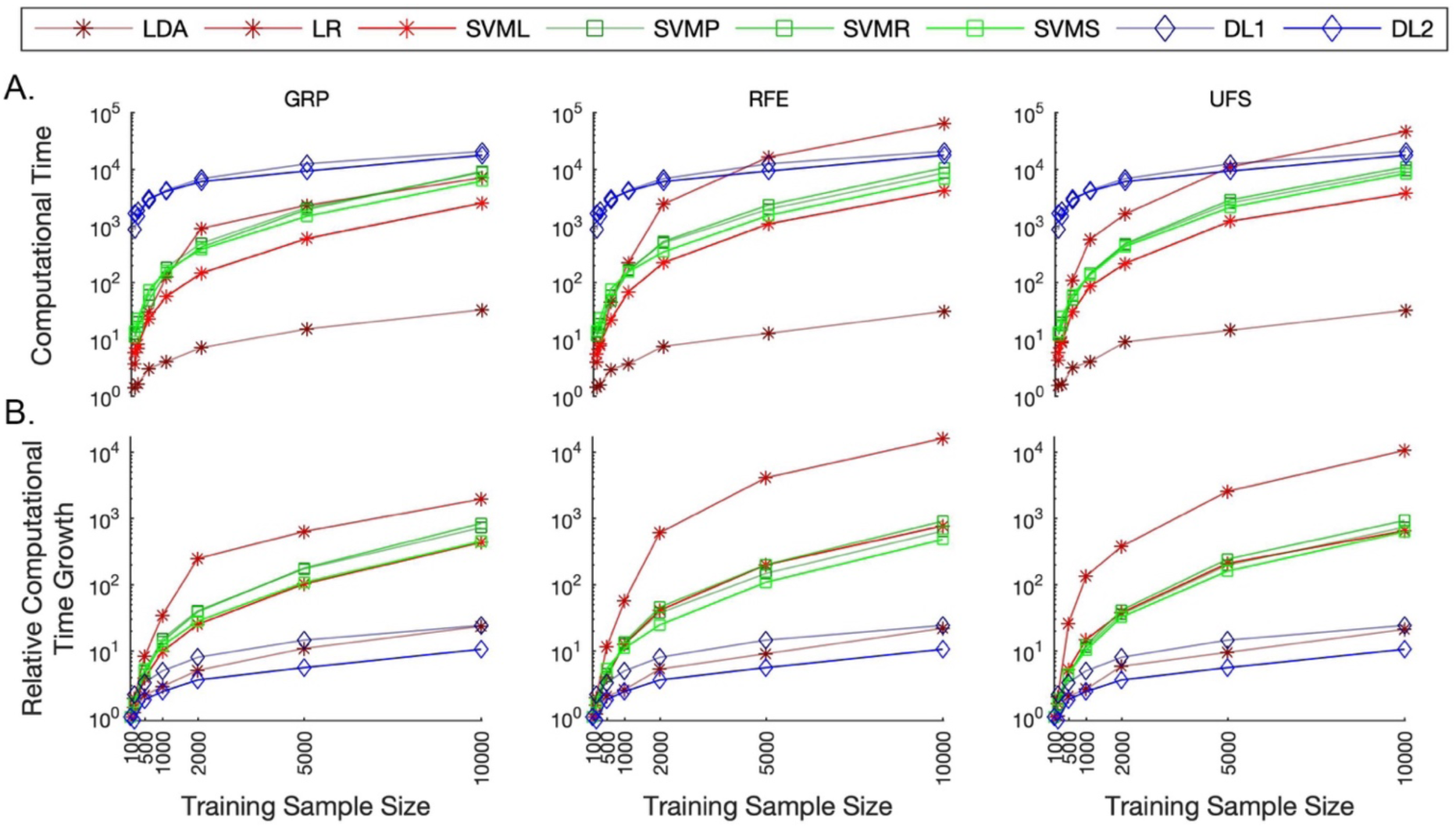
Systematic Comparison of Computational Time Complexity. This analysis compared the growth in computational time with an increase in the training sample size across all models and dimension reduction methods. **(A)** The DL models recorded higher run times but showed a more linearly increasing trend with an increase in training sample size. On the other hand, the SML models presented a quadratic trend (except for the LDA method). As a result, the difference between the recorded run times between the two classes of models decreased as the training sample size increased. **(B)** To further validate this trend, we conducted a similar analysis on a metric for relative computational time growth (defined as the computational time for the given training sample size normalized by this metric for the smallest sample size). This analysis confirmed a lower asymptotic complexity in the relative run time growth for DL models.

Furthermore, we note that the computation time of the SML models did not include the time used for the dimensionality reduction. Additionally, the GPU implementation for DL models used the same number of CPU threads (*n* = 8) as that for the SML models. As the relative computational time metric was estimated relative to a baseline (i.e. smallest sample size for the same method), we speculate the difference in nature of the implementations for the two classes of methods to not result in a significant change in the relative metric, even though the non-normalized metric in **Figure 2A** could be expected to drop for the SML models if a GPU based implementation were used. Critically, our focus on comparing the computational growth rate for the commonly used modern implementations of these models assumes high significance as bigger data samples are becoming increasingly available for research and can be safely expected to grow even bigger in coming years.

### DL Learns Meaningful Brain Representations that Span a Comprehensible Projection Spectrum

If the DL methods are indeed learning embedding that is representing the brain in low dimensional space, the encodings in deeper layers (further from the input) must be discriminative for the attempted task. Thus, for the undertaken classification task in this work, we can expect the DL encodings to capture meaningful age and gender information from the high-dimensional input data. Furthermore, we can anticipate such information in the captured patterns to continually distill with an increase in training sample size. To validate this statement, we conducted a post hoc analysis by projecting the learnt DL1 embeddings (i.e. the output of the first fully-connected layer in the DL1 architecture) onto a two-dimensional space using t-distributed stochastic neighbor embedding (t-SNE) ^29^ for the entire range of training sample sizes, and color-coding the two-dimensional projection spectrum by the class labels. The t-SNE algorithm works on placing two dimensional representations maximally preserving their distances in the original space; thus, if the embeddings contain pronounced age and gender information, subjects of the same gender and similar age are expected to end up nearby. The t-SNE layouts of the learnt DL representations in **Figure 3** reveal meaningful refinement of the learnt patterns with increasing training sample size, with the progressive evolution of an explicit bi-modal structure (i.e. formation of two distinct gender clusters) both modes of which manifest a comprehensible, gradual spectrum of age. More specifically, we can see separate gender clusters ordered in increasing age from one end of the spectrum to the other, although some outlier observations do exist. Hence, we can conclude that the implemented method was indeed able to learn the representational patterns of interest from brain imaging data.

**Figure 3:**
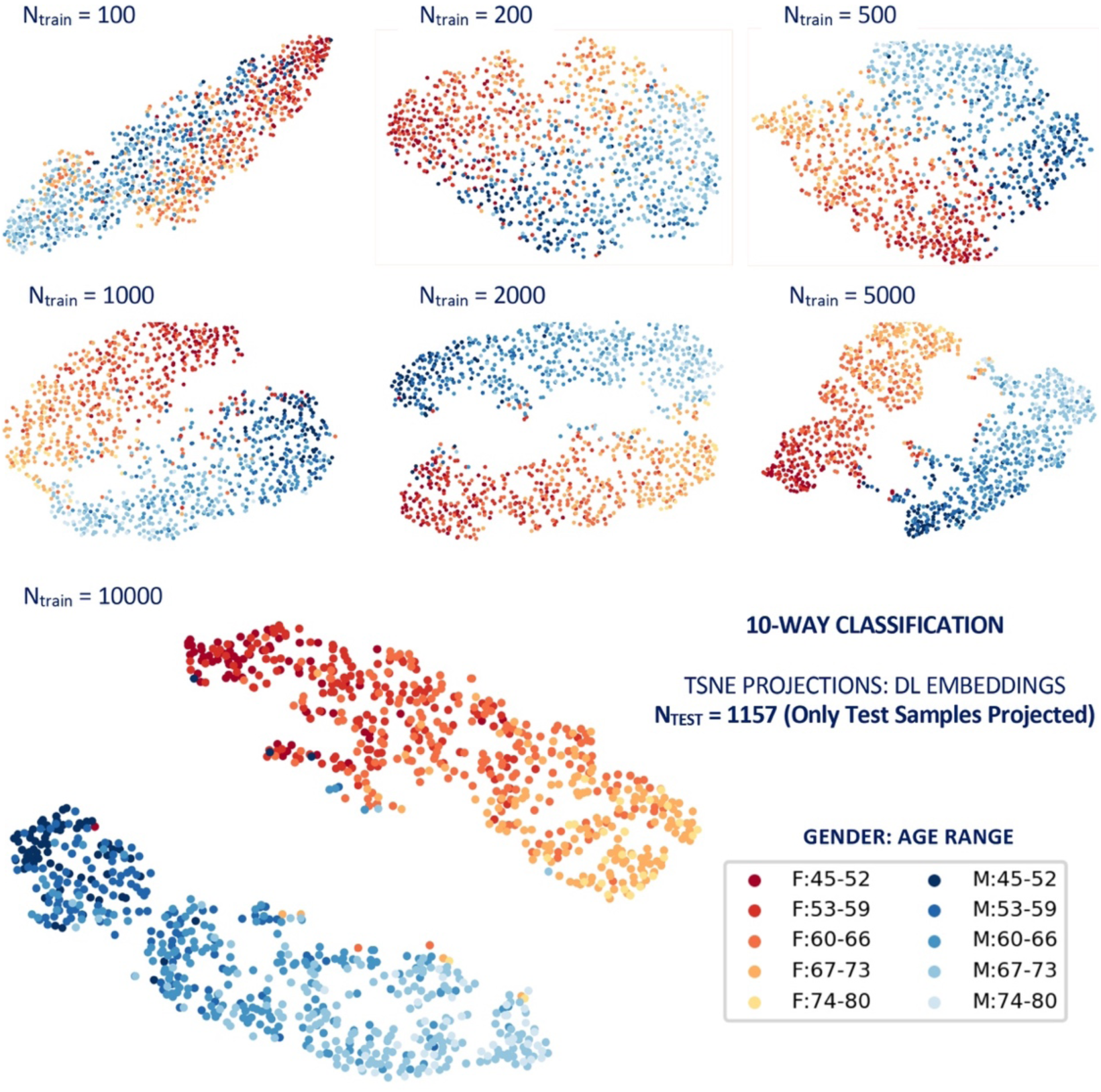
Projections of the embeddings from the validated DL models compared across a range of training sample sizes. Representational patterns of the brain are indeed learnt. In fact, they distill continually with increasing training sample size, and eventually evolve into separate gender clusters (i.e. red/F/female and blue/M/male clusters), both presenting a gradual spectrum of age (i.e. traceable light colored to dark colored).

### DL Enables Robust Prediction Relevance Estimates for Human Brain Regions

A critical dimension for validating the robustness of an algorithm is the similarity in the predictions across its independent repetitions. Hence, we sought to determine if the validated DL models estimated prediction relevance of the brain regions in the classification decisions in a consistent pattern across their independent runs. For this, we recorded the saliency maps for the multiple repetitions (*n* = 20) that varied in the randomly sampled training, validation and testing data for the highest training size (*n* = 10000). The saliency maps were estimated through two standard approaches, namely gradient-based backpropagation ^30^ (GBP) and network occlusion sensitivity analysis ^31^ (NOSA). The GBP approach computes the gradient of the class score with respect to the input image to determine the relevance of each pixel with respect to the classification decision. In the NOSA method, brain functional networks are occluded one at a time, their class probabilities are re-evaluated, and their relevance in the classification decisions is estimated proportional to the reduction in target class probability. Both of these methods are sensitive to the data and network architecture ^32^, thus perfectly fitting in the scope of the attempted classification task in this work.

**Figure 4** presents the prediction relevance percentages based on the highest sample size computed for these approaches on the Automated Anatomical Labelling (AAL) brain atlas ^33^. Despite some variation in the ranking orders of the merged brain networks, both saliency approaches estimated similar prediction levels for most of the brain networks. The most discriminative brain networks highlighted through these approaches included peak activations in the middle temporal gyrus, middle frontal gyrus, superior frontal gyrus, postcentral gyrus, inferior temporal gyrus, cerebellum crus 1, precentral gyrus, fusiform gyrus, superior temporal gyrus, superior medial frontal gyrus, insula, middle occipital gyrus, middle cingulum gyrus and calcarine gyrus regions. The mean relevance estimates for the AAL brain atlas for both approaches and scatterplot of these prediction metrics comparing the two approaches (*r* = 0.921) are illustrated in **Figure 5**. Overall, these initial results clearly suggest robustness in the prediction relevance estimates and thus the high potential of the undertaken DL approach to record consistent representations of the brain imaging data. In view of such positive evidence, future DL applied to brain imaging data should investigate incorporate saliency mapping into learning formulations more comprehensively.

**Figure 4:**
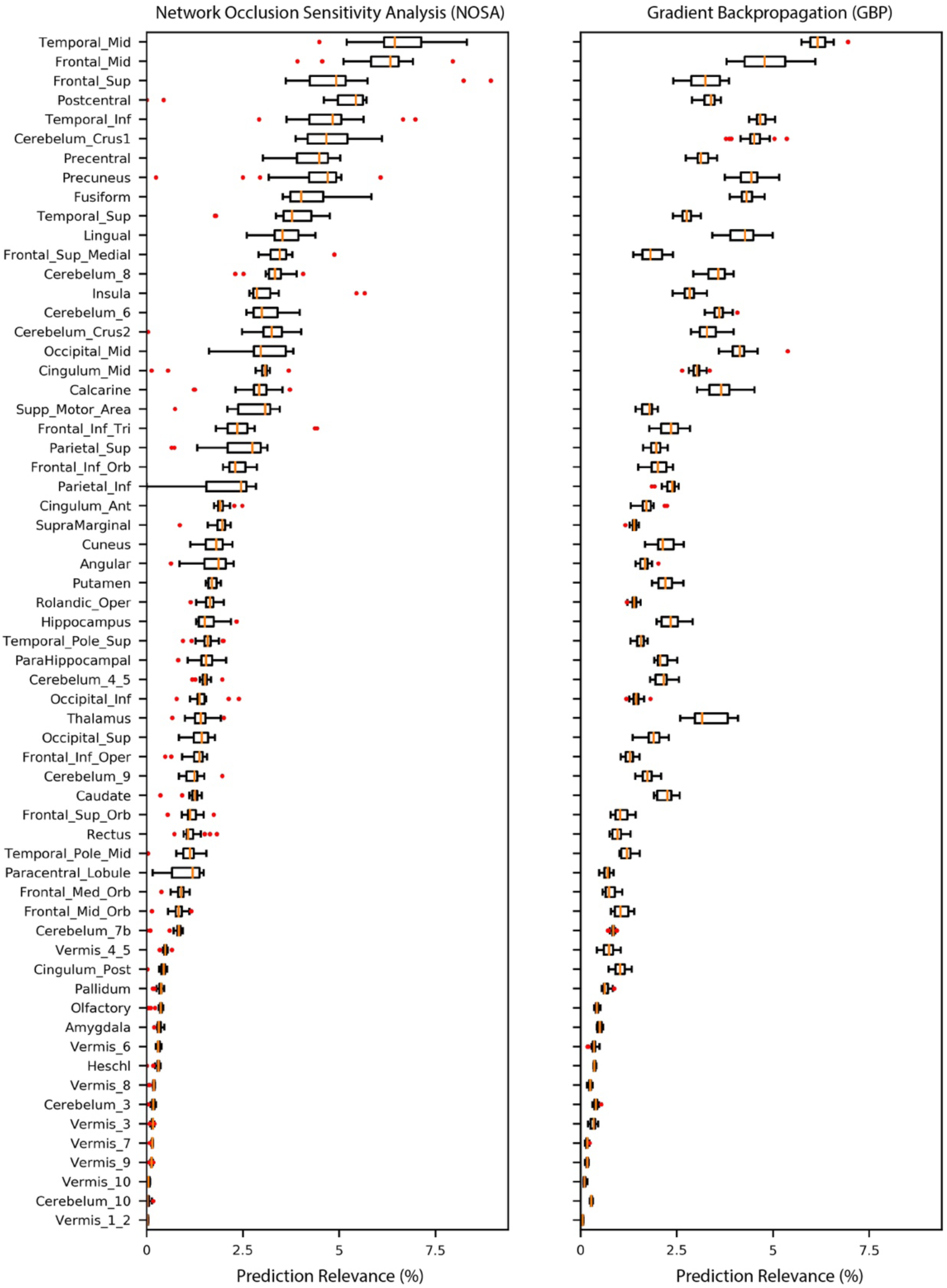
Prediction relevance estimates for the AAL atlas for the network occlusion sensitivity analysis (NOSA) and gradient backpropagation (GBP) approaches. Each boxplot outlines the variation in the prediction relevance percentages for the corresponding brain region across twenty independent runs of the classification task. These estimates generally spanned a narrow range (except for few outliers runs for some brain regions), and a comparison of these standard approaches confirmed consistency in the trends for most of the brain regions (see Figure 5 for clearer trends). Note, the AAL atlas brain regions are sorted from higher prediction relevance to lower prediction relevance for the NOSA approach for this illustration.

**Figure 5:**
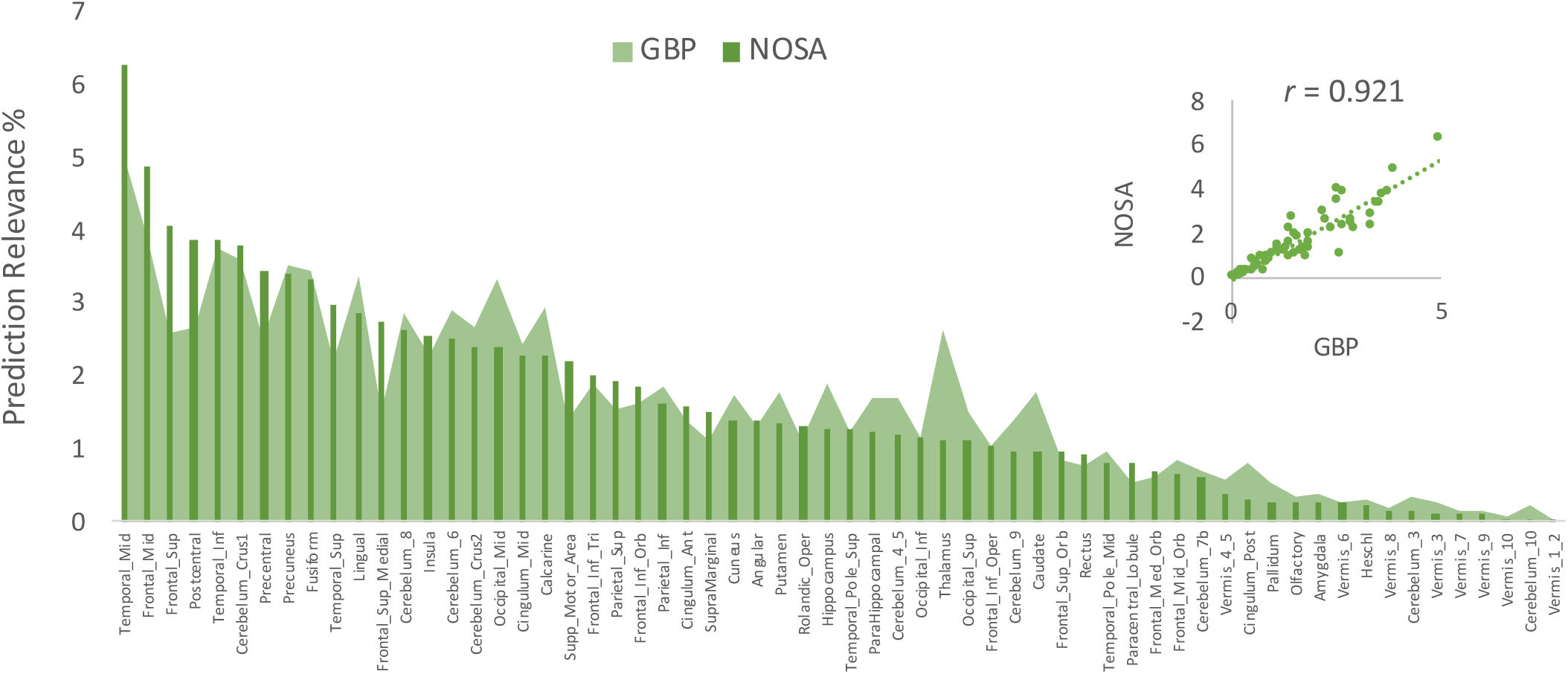
Mean prediction relevance estimates for the AAL atlas. The two tested saliency mapping methods: Gradient Backpropagation (GBP) and Network Occlusion Sensitivity Analysis (NOSA) show a high correlation value of 0.92 in the mean prediction relevance estimates, thus confirming the consistency in the prediction relevance from the learned DL representations.

## Discussion

Our results demonstrate that DL methods, if implemented and trained according to the common practices, have the potential to substantially outperform SML methods and to scale particularly well presenting a lower asymptotic complexity in relative computational time, despite being more complex in their architecture and parameterization. This observation of daylight margins in the performance graphs is consistent with the notion that the studied task associations are embedded at intricate abstract levels in the complex imaging data and therefore can benefit from the representational power of the DL methods. We further corroborated this notion by demonstrating that superior feature extraction contributed to the excellent performance of DL methods and that SML methods can perform equally well if we train them on the DL representations. Hence, we strongly recommend future work with DL methods to assess the wealth of spatiotemporal information available in the minimally preprocessed input space, as compared to working with reduced feature spaces. Note, we are not discovering something new here, the DL field is not only aware of this property of the models, but arguably they were developed with this as a primary goal ^5,6,34,35^. Our analysis also suggests that the performance improvement as a function of training sample size for DL methods eventually saturates similarly to SML methods, although at a significantly higher performance mark. Although the deeper variant of the DL method tested in this work trained faster, it did not result in a significantly improved performance, therefore demanding a further probe to confirm if additional depth could further enhance the performance of the DL models. We note here that there are, nevertheless, many gateways to potentially score further performance gains apart from experimenting with even deeper variants of the tested class of DL models, for example, exploring variations in the finetuning process and testing other existing or newer DL frameworks. Indeed, it would be highly interesting to benchmark the performance and scalability bounds of an extensive battery of diverse supervised and unsupervised DL frameworks on brain imaging data.

We also illustrated that the DL embeddings from the validated models indeed capture meaningful and consistent patterns of representations of the brain that distilled continuously with more training data, spanning a comprehensible projection spectrum of age and gender. We further exemplified this by showing that DL consistently localizes discriminative brain representations, demonstrating the robustness of prediction relevance estimates in multiple relevance interpretation methods. These latter results reveal the high potential of DL methods in learning specific changes in the brain that explain the differences in the analyzed groups, an attribute that holds paramount significance in important applications such as disease characterization and studying treatment effects. We presume the effectiveness of engaging DL methods in the domain of brain imaging would primarily depend on how well we can explain such vital tasks, by leveraging the existing as well as by building newer methodical interpretations for the diverse range of DL models. Eventually, the DL chalk-horses in neuroimaging would highly likely be the ones that source a healthy blend of superior representational learning and finer interpretability.

In essence, our findings highlight the presence of non-linearities in the brain imaging data that DL frameworks can exploit to generate more predictive encodings for characterizing the human brain. Results are in support of the potential of DL applications to brain imaging data, even with currently available data sizes; existing claims/speculations of the unlimited scalability of DL methods, however, demand further confirmation. Our findings motivate future DL work in brain imaging to focus on excelling in the predictive power of the encodings and facilitating more precise discriminative feature localization through methodical model interpretations. Notably, the predictive capacity of DL models is more straightforward to evaluate, but that is not the only and, arguably, the primary use that can benefit from them. A number of other applications such as segmentation and multimodal data integration directly benefit from representational powers and ease of model construction of the DL approaches. Rather than focusing on ways to show DL does not predict as well in some instances, we should be leveraging the flexibility of these models, to greatly advance in brain imaging problems that the current workhorse SML methods are not able to further push.

## Methods

### Data

This work used sMRI images (*n* = 12,314) from unaffected subjects (i.e. those who had no diagnosed or self-reported mental illnesses based on 22,392 subjects’ sMRI data available as of April 7, 2019) from the UK Biobank repository. The sMRI data were segmented into gray matter, white matter and cerebral spinal fluid. The gray matter images were then warped to standard space, modulated and smoothed using a Gaussian kernel with an FWHM = 10 mm. The preprocessed gray matter volume images had a dimensionality of 121 x 145 x 121 in the voxel space, with the voxel size of 1.5 x 1.5 x 1.5 mm^3^.

### Cross-validation Procedure

A key objective of this work was to compare how classification performance on the human brain images scaled with increasing sample sizes across the different SML and DL models. To execute this, the dataset of 12314 subjects was stratified into three partitions: training (*n* = 10000), validation (*n* = 1157) and test (*n* = 1157), with the training sample size varying in the range 100 to 10000 subjects (*n* = 100, 200, 500, 1000, 2000, 5000 and 10000). For all tested (SML and DL) models, a repeated (*n* = 20) stratified monte-carlo (i.e. repeated random sub-sampling) cross-validation (CV) procedure was employed. For each repetition, the training, validation and test samples were sampled exactly once to ensure a consistent comparison by keeping them the same across the different methods. Notably, the above procedure was repeated to decrease the estimator bias and generate better estimates of the distribution of the classification performance metric. Finally, hyper-parameter tuning (detailed in a following section) was employed using the training and validation folds and held-out test data samples were fed to the validated models to compute test accuracies for each CV repetition and classification approach. In summary, the primary performance comparison was employed across multiple dimensions: twenty repetitions, seven training sample sizes, three/none (SML/DL) dimension reduction methods and six/two (SML/DL) classifiers.

### SML Models

Six linear and non-linear SML models (motivated by Schulz, et al. ^26^) were tested to estimate a diverse baseline to compare the performance of DL models. The linear approaches included linear discriminant analysis (LDA) method, logistic regression (LR) and support vector machines with a linear kernel (SVML) models, whereas the non-linear SML models included support vector machines with a polynomial (SVMP), radial-basis (SVMR) and sigmoidal kernel (SVMS).

### Feature extraction for SML models

Feature extraction is a crucial process to eliminate redundant features in the data to boost the performance of linear and kernel-based algorithms ^3^. This step helps in reducing overfitting as lesser data dimensions imply a lesser probability of inferring decisions based on noise, and also significantly reduces the model complexity and training time of the algorithms. Hence, three dimensionality reduction methods (DRMs) were tested for feature extraction (as in Schulz, et al. ^26^) for all six SML models: Gaussian random projection (GRP), recursive feature elimination (RFE) and univariate feature selection (UFS). The GRP method projects the high-dimensional data onto a lower-dimensional subspace preserving the similarity in the data vectors using a random matrix generated using a Gaussian distribution ^36,37^. Recursive feature elimination is an iterative procedure that prunes the least ranked (i.e. least significant) features in each iteration until the desired number of features is achieved ^38,39^. In this analysis, we reduced 25% of the least ranked features at each step. Finally, the UFS method is based on univariate statistical tests to select the highest scoring features. We used the ANOVA F-values to return the univariate scores for feature ranking. For a consistent comparison and following the same work, the (voxel-wise) input space for each subject was reduced to a 784-dimensional subspace using each of these methods.

### Hyperparameter Validation

Hyperparameter tuning was employed for SML models through a grid parameter search with the *hypopt* python package. The hyperparameter grids were chosen to be consistent with the commonly evaluated parameter ranges as also employed in Schulz, et al. ^26^. The cost parameter (that is proportional to the inverse of the regularization strength) was tested for ten values sampled on a logarithm scale for a range of powers of 2 from [-20, 10] for the LR and SVML models. This range of powers was shifted for all three non-linear SVM kernel models to [-10, 20]. The other regularization parameter, gamma, was tested for 10 values sampled on a logarithm scale for a range of powers of 2 from [-25, 5] for all non-linear SVMs. Coefficients for the poly and sigmoid kernel SVMs were tested for -1, 0 and 1 values. To ensure computational tractability, the maximum number of iterations was set to 10000 for all four SVM models. The number of CPU threads to be used for the grid search class in the hypopt package was set to 8, and this parameter was kept the same in the DL training routines to allow a consistent comparison in the empirical asymptotic relative time complexity.

### DL Models

Two 3D-CNN variants of the AlexNet architecture ^28^ were implemented on the open-source Pytorch GPU framework to establish a performance baseline for the DL models (Figure 6). The first variant (DL1) was configured with five convolutional layers with a variable number of channels in each of the convolutional layers (64C-128C-192C-192C-128C). In contrast, the second variant (DL2) was a *deeper* version with six convolutional layers and an increased number of channels in the later layers (64C-128C-192C-384C-256C-256C). Detailed specifications of the 3D convolutional, 3D batch normalization, non-linear activation (ReLu), 3D MaxPooling, Dropout and Linear (i.e. fully connected) layers for both DL architectures are illustrated in Figure 6.

**Figure 6:**
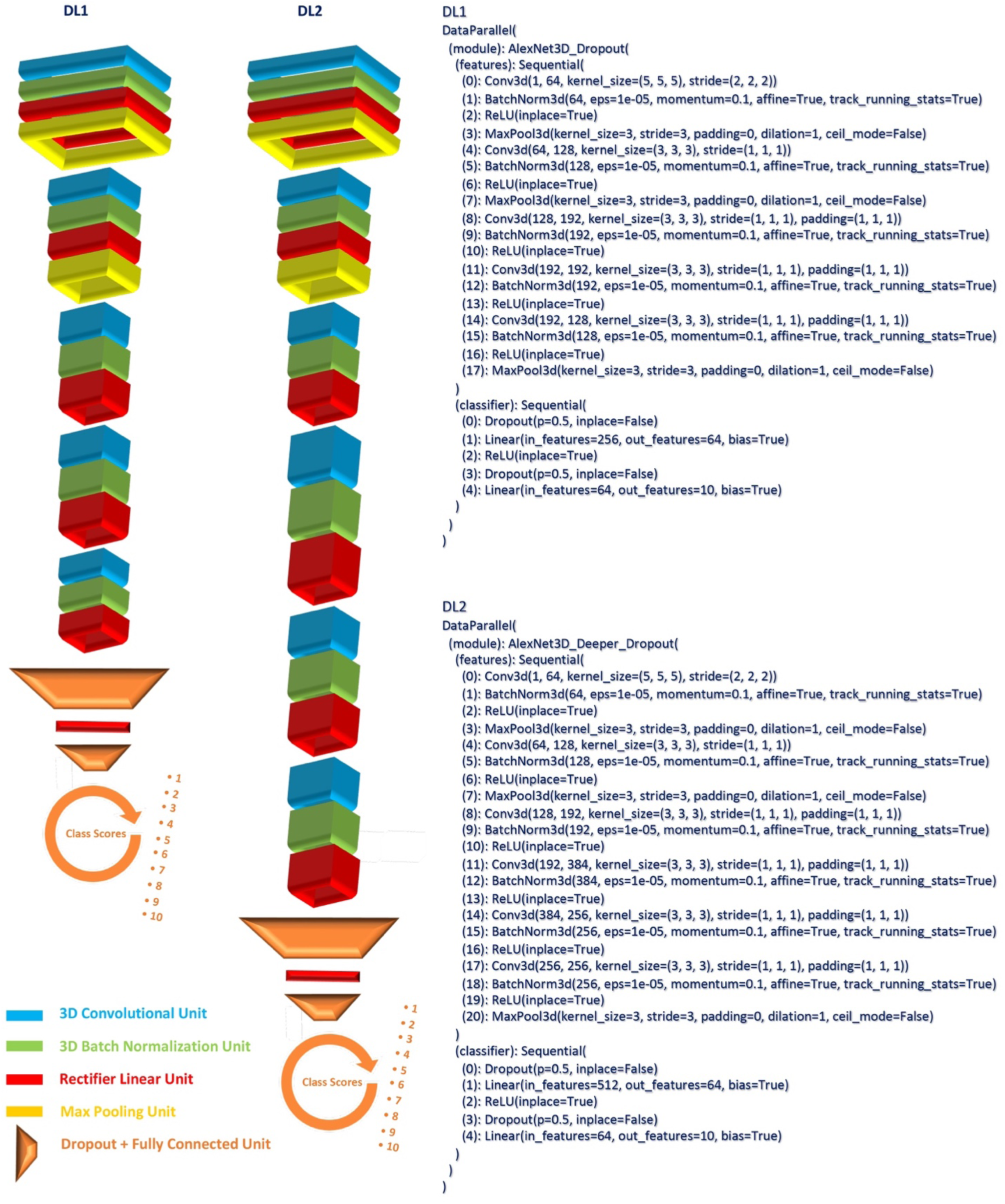
High level architecture visualization and detailed specification of the two deep learning models used in this work.

### DL training

Training and testing routines for the DL architectures were implemented on an NVIDIA CUDA parallel computing platform (accessing 2 Intel(R) Xeon(R) Gold 6230 CPU @ 2.10GHz nodes on the TReNDS cluster each with 4 NVIDIA Tesla V100 SXM2 32 GB GPUs) using GPU accelerated NVIDIA CUDA toolkit and Pytorch tensor libraries. The Adam algorithm ^40^ for first-order gradient-based optimization of stochastic objective functions was preferred for its computational efficiency, low memory requirements, and suitability for tasks with high-dimensional parameter spaces. For the DL1 method, a batch size of 32 and learning rate parameter of 1×10−5 was validated in the hyperparameter tuning stage from an initial grid parameter search (batch size: [2, 4, 8, 16, 32, 64] and learning rate: [1×10−1, 1×10−2, 1×10−3, 1×10−4, 1×10−5, 1×10−6]] on a randomly chosen largest training sample size crossvalidation fold. These parameters were retained for the second DL architecture due to the high similarity in the architectures. A learning rate scheduler callback was employed to reduce the learning rate by a factor of 0.5 on plateauing of the validation accuracy metric. Early stopping with a patience level of 20 epochs was implemented on the validation accuracy metric to reduce overfitting and achieve lower generalization error in the testing phase. Similar to the SML models, a maximum number of 8 CPU threads were allocated to each of the DL runs for a consistent comparison in the time complexity analysis.

## Acknowledgements

This work was supported by National Institutes of Health (NIH) grants R01EB006841, RF1AG063153 and R01EB020407 to Dr. Vince D. Calhoun.

## Author Contributions

Conceptualization: A.A., S.P. and V.C.

Funding Acquisition: V.C.

Supervision: S.P. and V.C.

Data Collection and Preprocessing: Z.F., M.S., R.S. and Y.D.

Methodology: A.A.

Software: A.A.

Validation: A.A.

Visualization: A.A.

Writing – original draft: A.A.

Writing – revisions: A.A., S.P. and V.C.

All authors have given final approval of this version of the article.

## Data Availability

The MRI data used in this work are available via the UKB website (https://www.ukbiobank.ac.uk).

## Code Availability

The code and python state dictionaries of the validated deep learning models developed for this work will be made available at https://github.com/aabrol/SMLvsDL/.

